# A two-locus hybrid incompatibility is widespread, polymorphic, and active in natural populations of *Mimulus*

**DOI:** 10.1101/339986

**Authors:** Matthew P. Zuellig, Andrea L. Sweigart

## Abstract

Reproductive isolation, which is essential for the maintenance of species in sympatry, is often incomplete between closely related species. In these taxa, reproductive barriers must continue to evolve within species, without being degraded by ongoing gene flow. To better understand this dynamic, we investigated the frequency and distribution of incompatibility alleles at a two-locus, recessive-recessive hybrid lethality system between species of yellow monkeyflower *(Mimulus guttatus* and *M. nasutus)* that hybridize in nature. We found that *M. guttatus* typically carries hybrid lethality alleles at one locus *(hl13)* and *M. nasutus* typically carries hybrid lethality alleles at the other locus *(hl14).* As a result, most naturally formed hybrids will carry incompatible alleles at both loci, with the potential to express hybrid lethality in later generations. Despite this general pattern, we also discovered considerable polymorphism at both *hl13* and *hl14* within both *Mimulus* species. For *M. guttatus*, polymorphism at both loci even occurs within populations, meaning that incompatible allele pairings might also often arise through regular, intraspecific gene flow. By examining genetic variation linked to *hl13* and *hl14,* we discovered that introgression from *M. nasutus* is a primary driver of this polymorphism within *M. guttatus.* Additionally, patterns of introgression at the two hybrid lethality loci suggest that natural selection acts to eliminate incompatible allele pairings, providing evidence that even weak reproductive barriers might promote genomic divergence between species.

## INTRODUCTION

The evolution of reproductive barriers is essential for the establishment and maintenance of species that co-occur in sympatry. Hybrid incompatibilities that manifest as hybrid sterility or inviability are common contributors to reproductive isolation in many species pairs (Coyne and Orr 2004) and recent studies have identified the loci, and in some cases the genes, that cause hybrid dysfunction (reviewed in Presgraves 2010, Maheshwari and Barbash 2011, Sweigart and Willis 2012, Fishman and Sweigart 2018). This work has provided novel insights into the molecular functions of hybrid incompatibility alleles and provided some hints about their evolutionary histories. However, we know little about the dynamics of hybrid incompatibilities between naturally hybridizing species, even though it is in such populations that we might directly assess their relevance to speciation. Do hybrid incompatibilities affect rates of gene flow in natural populations? What evolutionary forces affect the frequency of incompatibility alleles within species? How are incompatibilities maintained among species that that experience ongoing interspecific gene flow?

A natural starting point for determining whether hybrid incompatibilities affect rates of contemporary interspecific gene flow is to assess their distribution and frequency within species, which can indicate whether incompatibilities are expressed in natural hybrids. In diverse taxa, incompatibility loci are often polymorphic *within* species, with both incompatible and compatible alleles existing at a single locus (e.g., Reed and Markow 2004, Shuker et al. 2005, Sweigart et al. 2007, Seidel et al. 2008, Good et al. 2008, Cutter 2012, Sicard et al. 2015, Larson et al. 2018). Such polymorphism may exist because of evolutionary processes acting within species, such as balancing selection maintaining multiple alleles (Seidel et al. 2008, Sicard et al. 2015) or incomplete selective sweeps (Sweigart and Flagel 2015). Alternatively, polymorphism might result from interspecific gene flow, with incompatible alleles being replaced by introgression of compatible alleles. This second possibility is expected to arise when hybrid incompatibilities are actively expressed in nature. Under such a scenario, theory predicts that incompatible alleles will be maintained at migration-selection balance or actively purged from the species to reduce the negative fitness consequences of hybridization (Gavrilets 1997, Kondrashov 2003, Lemmon and Kirkpatrick 2006, Feder and Nosil 2009, Bank et al. 2012). Thus, characterizing hybrid incompatibility alleles that occur among naturally hybridizing species has the potential to reveal important dynamics that affect the long-term maintenance of incompatibilities in nature.

In this study, we investigate the frequency and distribution of incompatibility alleles that cause lethality in the hybrid progeny of yellow monkeyflowers, *Mimulus guttatus* and *M. nasutus.* These species are ideal candidates for studying the evolutionary dynamics of hybrid incompatibilities in nature, because they often occur in secondary sympatry and exhibit patterns of both contemporary and historical introgression, largely caused by unidirectional gene flow from *M. nasutus* to *M. guttatus* (Sweigart and Willis 2003, Brandvain et al. 2014, Kenney and Sweigart 2016). We recently discovered a genetically simple hybrid incompatibility that occurs between two inbred lines of *M. guttatus* (DPR102-gutt) and *M. nasutus* (DPR104-nas) derived from the sympatric Don Pedro Reservoir (DPR) population in central California (Zuellig and Sweigart 2018). In that study, we showed that one sixteenth of reciprocal F2 hybrids die in the cotyledon stage of development due to a complete lack of chlorophyll production. Lethality occurs in F2 hybrids that are homozygous for DPR102-gutt alleles at the *hybrid lethal 13 (hl13)* locus and homozygous for DPR104-nas alleles at the *hybrid lethal 14 (hl14)* locus. We also uncovered the molecular genetic basis of this two-locus incompatibility: hybrids lack a functional copy of the essential photosynthetic gene *PLASTID TRANSCRIPTIONALLY ACTIVE CHROMOSOME 14 (pTAC14),* which was duplicated from *hl13* to *hl14* in *M. guttatus* but not in *M. nasutus* (Figure 1A). Because hybrid lethality is caused by a non-functional copy of *pTAC14* at *hl13* in *M. guttatus* combined with the lack of *pTAC14* at *hl14* in *M. nasutus,* it seems reasonable to assume that these incompatibility alleles are selectively neutral *(i.e.,* natural selection is not acting to maintain incompatible alleles within either species; Muller 1942, Werth and Windham 1991, Lynch and Force 2000). Importantly, neutral incompatibility alleles are expected to be purged from species undergoing even low levels of ongoing gene flow and signatures of introgression of compatible alleles should be apparent in the genome (Gavrilets 1997, Kondrashov 2003, Lemmon and Kirkpatrick 2006, Feder and Nosil 2009, Bank et al. 2012, Muir and Hahn 2014).

**Figure 1:**
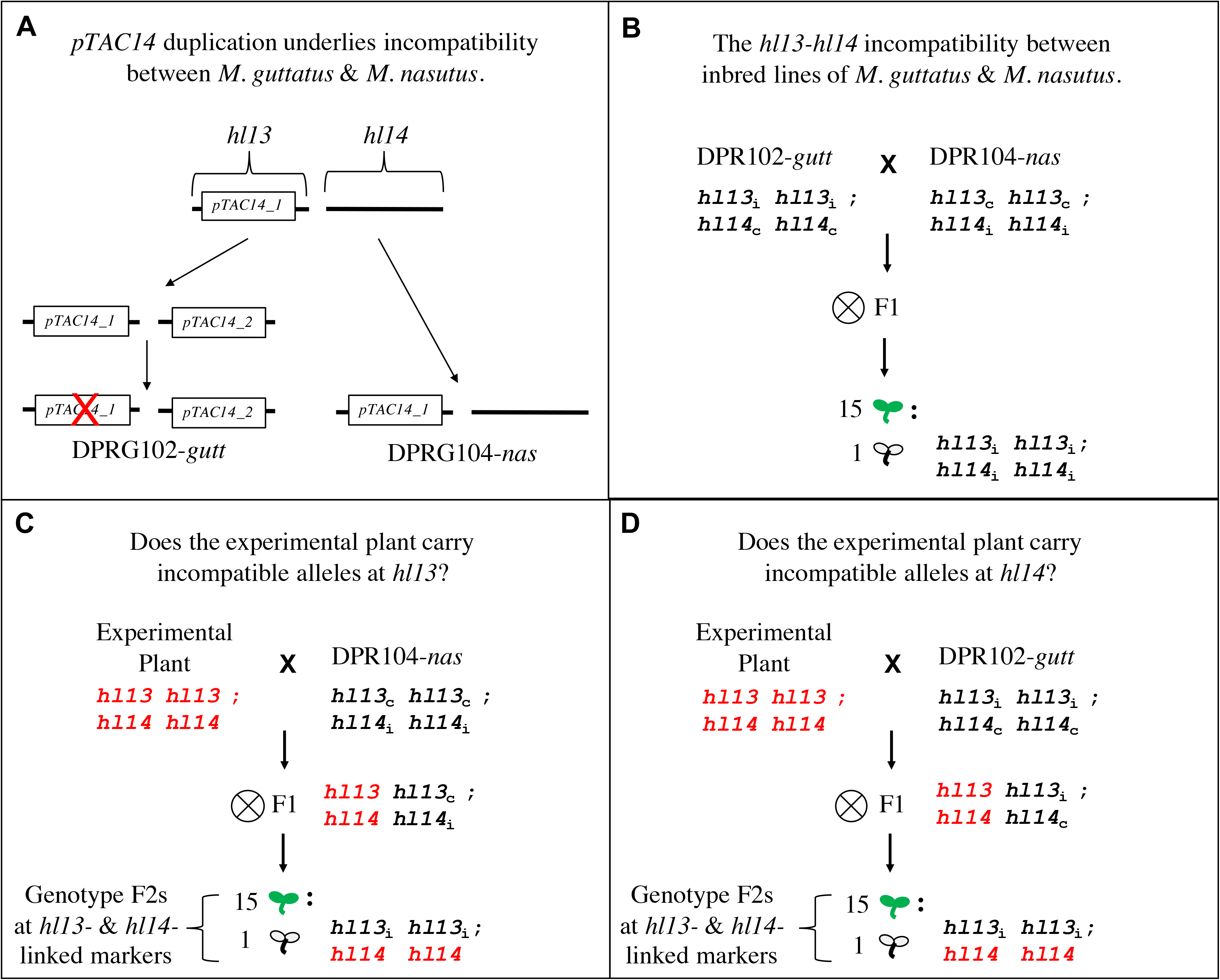
Crossing design used to characterize variation for the *hl13lhl14* incompatibility in experimental individuals. A) Hybrid lethality between inbred lines of *M. guttatus* (DPR10-gutt) and *M. nasutus* (DPR104-nas) is caused by the duplication and subsequent loss-of-function (red X) of ancestral *pTAC14* in *M. guttatus.* One sixteenth of F2 progeny will therefore carry nonfunctional *pTAC14* at *hl13* (from *M. guttatus)* and no copy at *hl14* (from *M. nasutus).* B) Illustration of the *hl13lhl14* incompatibility in the DPR102-gutt x DPR104-nas cross. DPR102-*gutt* is homozygous for incompatible ‘i’ alleles at *hl13* and compatible ‘c’ alleles at *hl14,* whereas DPR104-nas carries the opposite allelic combination. White F2 seedlings from this cross are homozygous for incompatible *hl13* alleles from DPR102-gutt *(hl13i)* and incompatible *hl14* alleles from DPR104-nas *(hl14ì).* C) Crossing design to determine whether experimental plants carried incompatible *hl13* alleles. Experimental plants were used as maternal parents in crosses to the tester DPR104-nas. From a single F1 hybrid, we generated F2 progeny, which we screened for the presence of white seedlings and genotyped for *hl13-* and hl14-linked markers. If white seedlings were homozygous for experimental plant alleles (red font) at *hl13* and homozygous for DPR104-nas alleles at *hl14,* we concluded that the experimental plant carried an incompatible allele at *hl13.* D) Crossing design to determine whether experimental plants carried incompatible *hl14* alleles. Experimental plants were used as maternal parents in crosses to the tester DPR102-gutt. From a single F1 hybrid, we generated F2 progeny, which we screened for the presence of white seedlings genotyped for *hl13-* and hl14-linked markers. If white seedlings were homozygous for experimental plant alleles (red font) at *hl14* and homozygous for DPR102-*gutt* alleles at *hl13*, we concluded that the experimental plant carried an incompatible allele at *hl14.*

Although the genes underlying hybrid lethality in *Mimulus* are now known (Zuellig and Sweigart 2018), we do not yet have a straightforward, sequencing-based approach to screen individuals for different *pTAC14* variants (duplicate copies of *pTAC14* are highly similar and distinguishing between them requires laborious cloning experiments). As an alternative, here we take a genetic crossing approach, generating F2 progeny between wild-collected plants and tester lines that carry known genotypes at *hl13* and *hl14.* Screening these F2 progeny for hybrid lethal (white) seedlings and genotyping them at *hl13*- and *hl14*-linked markers allows us to determine the incompatibility status of a large collection of *hl13-* and hl14-alleles from *M. guttatus* and *M. nasutus* sampled from throughout the species’ ranges. With these data, we estimate the frequency of incompatibility alleles within and among populations of each *Mimulus* species (even without knowing the underlying molecular lesions at the *pTAC14* duplicates). We also use the genotypes from *hl13-* and hl14-linked markers to begin investigating patterns of interspecific gene flow in genomic regions linked to hybrid lethality loci. As one of the most thorough studies of natural variation for a hybrid incompatibility to date, our findings provide unique insight into the evolutionary dynamics of incompatibility alleles and their role in reproductive isolation.

## METHODS

### *Mimulus* lines and crossing design

To investigate natural variation at *hl13* and *hl14* incompatibility loci, we intercrossed experimental plants from natural populations of *M. guttatus* and *M. nasutus* (Table S1) and “tester” lines with known *hl13-hl14* genotypes. Experimental plants came from two different sources: 1) wild-collected, outbred seeds from single fruits *(N* = 14), or 2) self-fertilized seeds of wild-collected plants (N = 64) (Figure 2, Table S1). For the primarily outcrossing *M. guttatus,* experimental plants generated from wild-collected seeds might carry as many as four alleles at each locus (or even more if there are multiple pollen donors), whereas experimental plants generated from the selfed progeny of a wild-collected plant should segregate only two alleles at each locus. For the highly self-fertilizing *M. nasutus*, experimental plants generated from either source are likely fixed for a single allele at each locus. As tester lines, we used previously generated inbred lines of *M. guttatus* (DPR102-gutt) and *M. nasutus* (DPR104-nas) (Zuellig and Sweigart 2018). DPR102-gutt is homozygous for incompatible alleles at *hl13* and compatible alleles at *hl14* (hl13_i_hl13_i_; *hl14_c_hl14_c_*), whereas DPR104-nas carries the opposite two-locus genotype (*hl13_c_hl13_c_*; *hl14_i_hl14_i_*) (Figure 1B).

**Figure 2:**
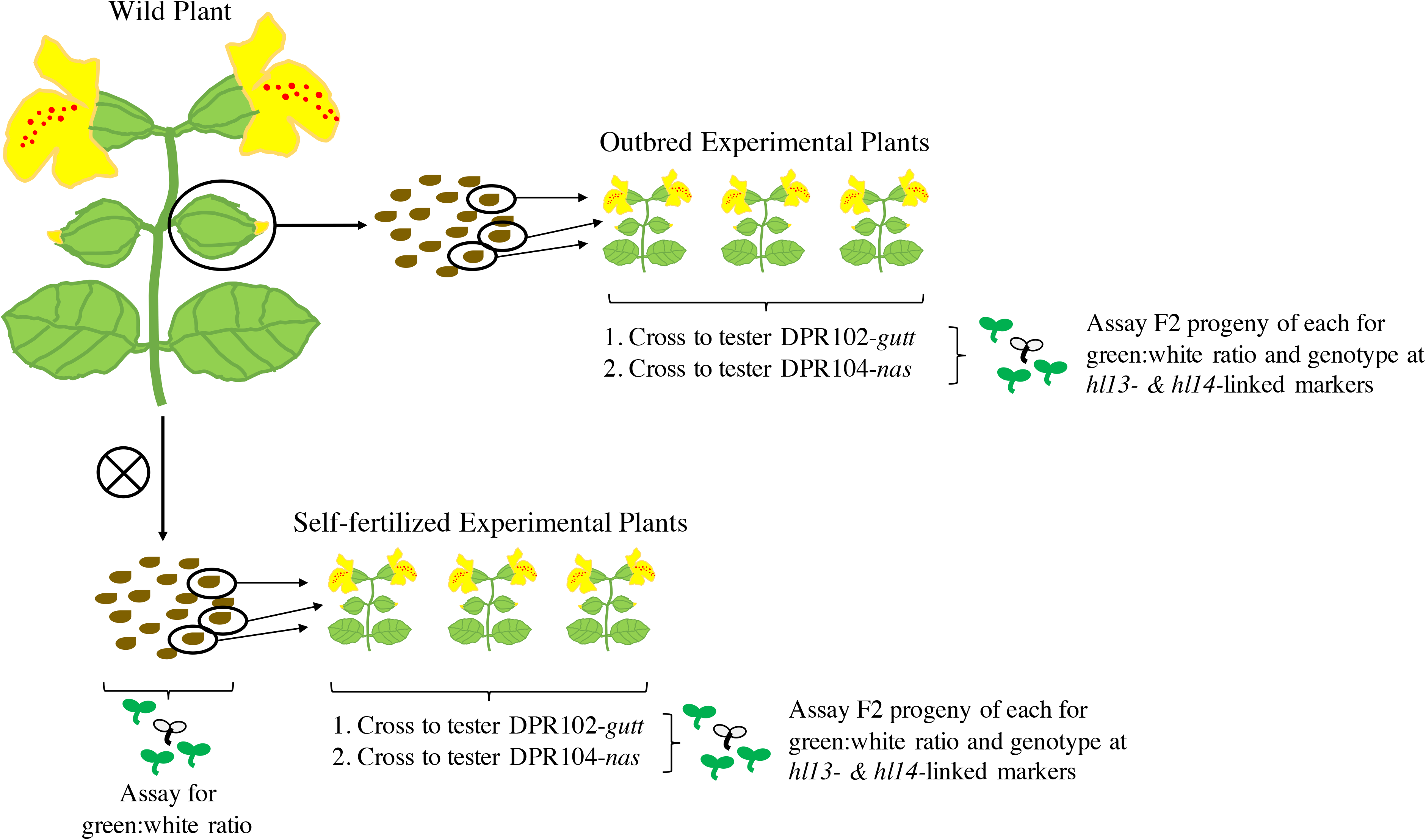
Sources of experimental plants used in genetic crosses. In some cases, “outbred experimental plants” were generated directly from wild-collected seeds. In other cases, wild plants were selfed to generate “self-fertilized experimental plants” and to assay seedlings for the chlorotic lethal phenotype. All experimental plants were crossed to DPR102-gutt and DPRN104-*nas* tester lines. From each cross, a single F1 hybrid was self-fertilized to generate F2 progeny, which were assayed for the presence of white seedlings *(i.e.,* green:white ratio) and genotyped at markers linked to *hl13* and *hl14.*

To test whether experimental plants carried incompatibility alleles at *hl13* or *hl14,* we generated F2 progeny for all experimental-tester crosses. We grew experimental plants to flowering and emasculated flowers prior to anther dehiscence. The following day, emasculated flowers were pollinated with one of the tester lines. We used a single F1 seed from each cross to generate an F2 progeny by self-fertilization (Figure 1C and D). By crossing experimental plants to the DPR102-gutt tester (which carries incompatible alleles at *hl13),* we were able to determine the incompatibility status of one *hl14* allele from the experimental parent *(i.e.,* whichever experimental plant allele was inherited by the F1 hybrid used to generate the F2). Similarly, by crossing experimental plants to the DPR104-nas tester (which carries incompatible alleles at *hl14),* we determined the *hl13* incompatibility status of one *hl13* allele from the experimental parent.

All plants used in this study were planted on moist Fafard-3B potting soil, cold stratified at 4°C for one week, and allowed to germinate in the University of Georgia greenhouses at 27°C under 16-hour days. Seeds were initially sown in 2.5” pots and thinned to a single plant after germination.

### Assessment of hybrid incompatibility status at *hl13* and *hl14*

For each of the 221 F2 progeny sets generated between experimental and tester plants (Table S2), we used a two-part strategy to determine the incompatibility status of the experimental parent’s allele at either *hl13* or *hl14.* First, we assayed phenotypes in the F2 progeny, surveying for the presence of white seedlings. In the original cross between DPR102-*gutt* and DPR104-nas, white seedlings occurred at a frequency of one-sixteenth in F2 hybrids (Figure 1, Zuellig and Sweigart 2018). In this study, if an F1 hybrid inherited an incompatible experimental allele at *either* hl13 or hl14, one of the two experimental-tester F2 progeny sets was expected to segregate white seedlings at a frequency of one sixteenth (Figure 1). If, on the other hand, an F1 hybrid inherited incompatible experimental alleles at *both* loci, the frequency of white seedlings in the F2 progeny was expected to be one-fourth. For each set of F2 progeny, we tallied the ratio of green to white for a minimum of 87 seedlings (average = 261, maximum = 751), which meant that, in every case, we had a >99% chance of observing white seedlings if the F1 hybrid carried an incompatible experimental allele at either locus (chance of observing at least one white seedling: 1-(15/16)^87^ = 0.996). For F2 progeny sets in which we saw at least one white seedling, the experimental allele (at either *hl13* or *hl14,* depending on the cross) was scored as potentially incompatible. For F2 progeny sets in which we observed only green seedlings, the experimental allele was scored as compatible at *hl13* or *hl14.*

The second step in our strategy to determine the incompatibility status of experimental alleles at *hl13* or *hl14* was to confirm that white seedlings in experimental-tester F2 progeny had inherited genotypes associated with the *hl13-hl14* incompatibility. In other words, we needed to ensure that hybrid lethality was due to *hl13* and *hl14,* and not to alleles at other genetic loci. For each wild plant, we determined *hl13-hl14* genotypes for at least one F2 progeny set containing white seedlings (we assumed that *hl13-hl14* incompatibility alleles identified in one experimental-tester F2 were also the cause of white seedlings in other F2 progeny descended from the same wild plant). For each of these F2 sets, we genotyped an average of 11 green (range = 1-33) and 13 white seedlings (range = 1-63) for size-polymorphic markers linked to *hl13* and *hl14.* We extracted genomic DNA from seedlings using a CTAB-chloroform protocol (Doyle 1987) modified for 96-well format. Using a standard touchdown PCR protocol, we amplified fluorescently-labeled markers linked to the hybrid sterility loci: M236 and M208 at *hl13,* and M132 and M241 at *hl14* (Figure S1). Although markers were originally designed to flank the hybrid sterility loci, we later discovered that both M208 and M236 are proximal to *hl13.* Because the closer of these two markers (M208) is only 134 kb (3.3 cM) from the causal *pTAC14* gene, its genotypes were used to infer *hl13* genotypes (genotyping error rate ~0.03 due to recombinants between M208 and hl13). To infer genotypes at *hl14,* we determined the genotypes of flanking markers M132 and M241 and excluded individuals with crossovers (based on the expected frequency of double crossovers, genotyping error rate ~0.01). This direct genetic approach allowed us to follow the inheritance of all *hl13*- and hl14-containing chromosomal fragments. All marker genotyping was performed by sizing PCR-amplified DNA fragments on an ABI3730XL automated DNA sequencer at the Georgia Genomics Facility. Marker genotypes were assigned automatically using GeneMarker (SoftGenetics LLC, State College, PA) and then verified by eye.

For each genotyped F2 progeny set, we scored the experimental parent’s allele as incompatible if seedling phenotypes were associated with hl13-hl14 genotypes (Figure 1). The presence of even a small number of white seedlings with the expected *hl13-hl14* genotype is strong evidence that the experimental parent carried an incompatible allele at *hl13* or *hl14* (probability of sampling one white seedling with a two-locus genotype consistent with the *hl13*-*hl14* incompatibility by random chance is 0.0625, probability of sampling two such white seedlings ~ 0.004; two or more white seedlings were genotyped in all but one F2).

### Determining frequencies of incompatibility alleles at *hl13* and *hl14* in natural populations

To determine *hl13* and *hl14* allele frequencies across the geographic ranges of *M. guttatus* and *M. nasutus*, we tallied the number of incompatibility alleles sampled in each population. It is important to note that because we often generated multiple experimental plants from a single wild individual (either from selfed or wild-collected seeds, Figure 2), alleles segregating in distinct sets of F2 progeny were not always independent *(i.e.,* the same wild allele might have been observed in more than one of its descendant F2s). Below we describe our methods for estimating the number of *hl13* and *hl14* alleles we sampled in wild individuals and populations.

For experimental plants generated from self-fertilized seeds (Figure 2), analysis of experimental-tester F2 progeny allowed us to directly infer the *hl13* and *hl14* genotypes of their wild parents (Tables S2, S3). We assumed that we sampled *both* of the wild plant’s alleles at a particular locus if we observed different phenotypes *(i.e.,* either one-sixteenth white seedlings or all green seedlings) in independent sets of descendant F2 progeny (indicating the wild plant was heterozygous), or if we observed the same phenotype in at least four sets of its descendant F2 progeny (indicating the wild plant was homozygous; probability of sampling only one wild allele in four sets of F2 progeny: 0.5^4^ × 100 = 6.25%). For cases in which the same phenotype was observed in three or fewer sets of F2 progeny descending from one wild plant, we assumed that we sampled only a single wild allele at that particular locus. In several cases, we obtained additional information about the wild plant’s *hl13-hl14* genotype by scoring the seedling phenotypes of its selfed progeny *(i.e.,* the siblings of experimental plants). For example, a lack of white seedlings in the selfed progeny often allowed us to infer that one of the two hybrid lethality loci must be homozygous for compatible alleles (see Table S3 for inferred genotypes).

For experimental plants generated from wild-collected, outbred seeds (Figure 2), alleles might have come from either the wild maternal plant or paternal pollen donor. Thus, unlike for experimental plants generated from selfed seeds, outbred experimental plants did not allow us to directly infer the *hl13-hl14* genotypes of the wild maternal parent (because wild alleles in the F2 might have descended from the pollen donor). Instead, for the 14 wild individuals that we assessed using outbred experimental plants, “inferred genotypes” (Table S3) are a composite of the alleles we found in their outbred progeny *(i.e.,* a snapshot of wild genotypes descended from that particular wild maternal plant). In most cases *(N* = 9), we used only a single outbred experimental plant per wild-collected fruit, effectively sampling one (maternal or paternal) allele from the wild (Table S2). However, for five wild plants (four *M. guttatus,* one *M. nasutus),* we generated two or three outbred experimental plants (Table S2). In these cases, the *hl13* and *hl14* alleles segregating in distinct sets of F2 progeny could not be considered independent, so we made the simple assumption that two of the four possible wild parental alleles were sampled (Table S3).

To calculate *hl13* and *hl14* allele frequencies in natural populations we inferred two-locus genotypes for 53 wild *M. guttatus* and 24 wild *M. nasutus* (using results from experimental-tester crosses and wild selfed progeny, Table S3). For each population, we determined the number of incompatible alleles per total alleles sampled.

### Assessment of linked variation at *hl13* and *hl14* in natural populations

Genotyping *hl13-* and hl14-linked markers in experimental-tester F2 hybrids allowed us not only to follow inheritance of the hybrid lethality alleles themselves, but also to investigate patterns of surrounding genomic variation. First, we determined marker allele sizes for the two tester lines, and then used this information to infer which alleles must have been inherited from the experimental parent. Thus, for both *hl13* and *hl14,* we were able to determine which marker variants occurred on chromosomes with compatible versus incompatible alleles (barring crossovers in the experimental parent inherited by the F1 hybrid, which would have disassociated M236-M208-hl13 alleles ~7% of the time and M132-hl14-M241 alleles ~11% of the time, Figure S1). For each hybrid lethality allele sampled, we determined its allele sizes at two linked markers (M236 and M208 for *hl13,* M132 and M241 for hl14), which we then combined into one composite “two-allele combination” (Tables S4 and S5). For both *hl13* and *hl14,* we used chisquare tests to determine whether linked marker allele combinations varied between species and/or hybrid lethality allele-type (compatible or incompatible).

## RESULTS

### Hybrid lethality alleles are widespread and polymorphic in *Mimulus*

To determine how often the *hl13/hl14* incompatibility occurs in nature, we investigated the distribution and frequency of hybrid lethality alleles in natural populations of *Mimulus* species from across their native ranges. By selfing wild-collected plants and crossing experimental plants to the DPR104-nas tester (which carries incompatible alleles at *hl14,* Figure 1), we determined the *hl13* incompatibility status of 58 *M. guttatus* alleles across 20 populations and 32 *M. nasutus* alleles across 10 populations (Tables S2 and S3). Similarly, by selfing wild plants and crossing experimental plants to the DPR102-gutt tester (which carries incompatible alleles at *hl13,* Figure 1), we determined the *hl14* incompatibility status of 68 *M. guttatus* alleles across 20 populations and 29 *M. nasutus* alleles across 11 populations (Tables S2 and S3). We discovered a pervasive role for the *hl13-hl14* incompatibility in chlorotic seedling mortality: across all F2 progeny of experimental and tester plants, nearly all white seedlings carried the incompatible two-locus genotype, whereas green seedlings carried the eight other two-locus combinations (in Mendelian ratios, *X^2^* = 2.560, *P* = 0.9225, df = 7) (Figure 3).

**Figure 3:**
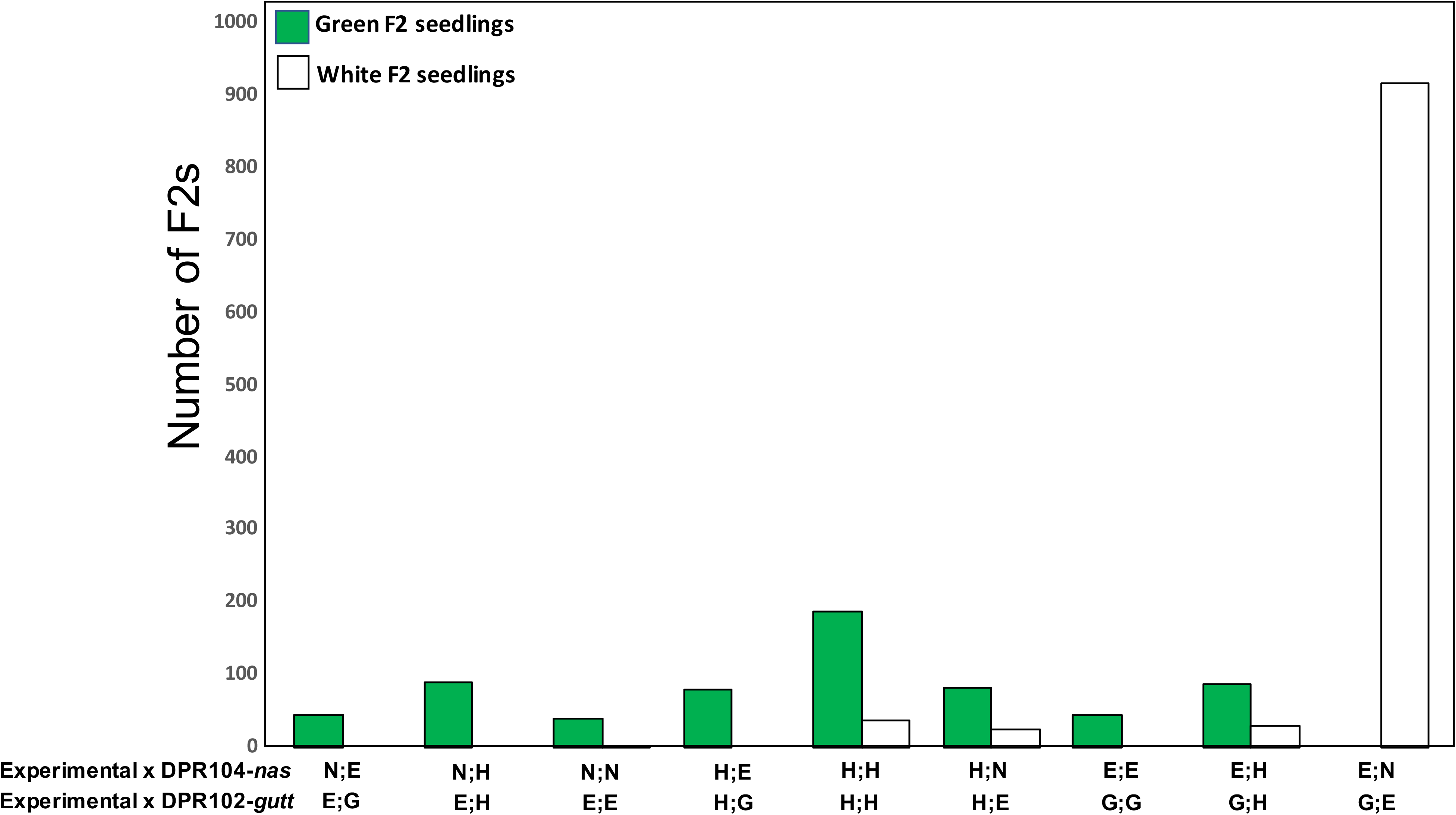
Frequency of two-locus *hl13-hl14* genotypes in green and white F2 seedlings. Green (N = 655) and white (N = 1012) seedlings from F2 families that segregated white seedlings are shown. White seedlings were oversampled in this study. Two-locus genotypes were inferred by genotyping seedlings at the same size-polymorphic markers linked to *hl13* and *hl14* that were used to determine linked marker allele combinations. The nine possible *hl13-hl14* two-locus genotypes are shown along the bottom for F2s generated when DPR102-gutt and DPR104-nas were used as paternal parents. “G” is homozygous for DPR102-gutt alleles, “N” is homozygous for DPR104-nas alleles, “E” is homozygous for experimental parent alleles, and “H” is heterozygous. Note that the “E;N” and “G;E” two-locus genotypes represent the expected incompatibility genotypes associated with the *hl13-hl14* incompatibility.

Remarkably, we discovered that both hybrid lethality loci are polymorphic within both *Mimulus* species. At *hl13,* the incompatibility allele is much more common in *M. guttatus* than in *M. nasutus*: it occurred at a frequency of 66% in *M. guttatus* and was found in all but one of the 20 populations surveyed, whereas it occurred at a frequency of 9% in *M. nasutus* and was found in 30% of populations (Figure 4A). At *hl14,* the situation is reversed with the incompatibility allele more common in *M. nasutus* than in *M. guttatus*: it occurred at a frequency of 62% in *M. nasutus* and was found in most (73%) of the populations surveyed, whereas it occurred at a frequency of only 25% in *M. guttatus* (Figure 4B). Despite this lower occurrence of the *hl14* incompatibility allele within *M. guttatus,* it was nevertheless sampled in 60% of the populations examined.

**Figure 4:**
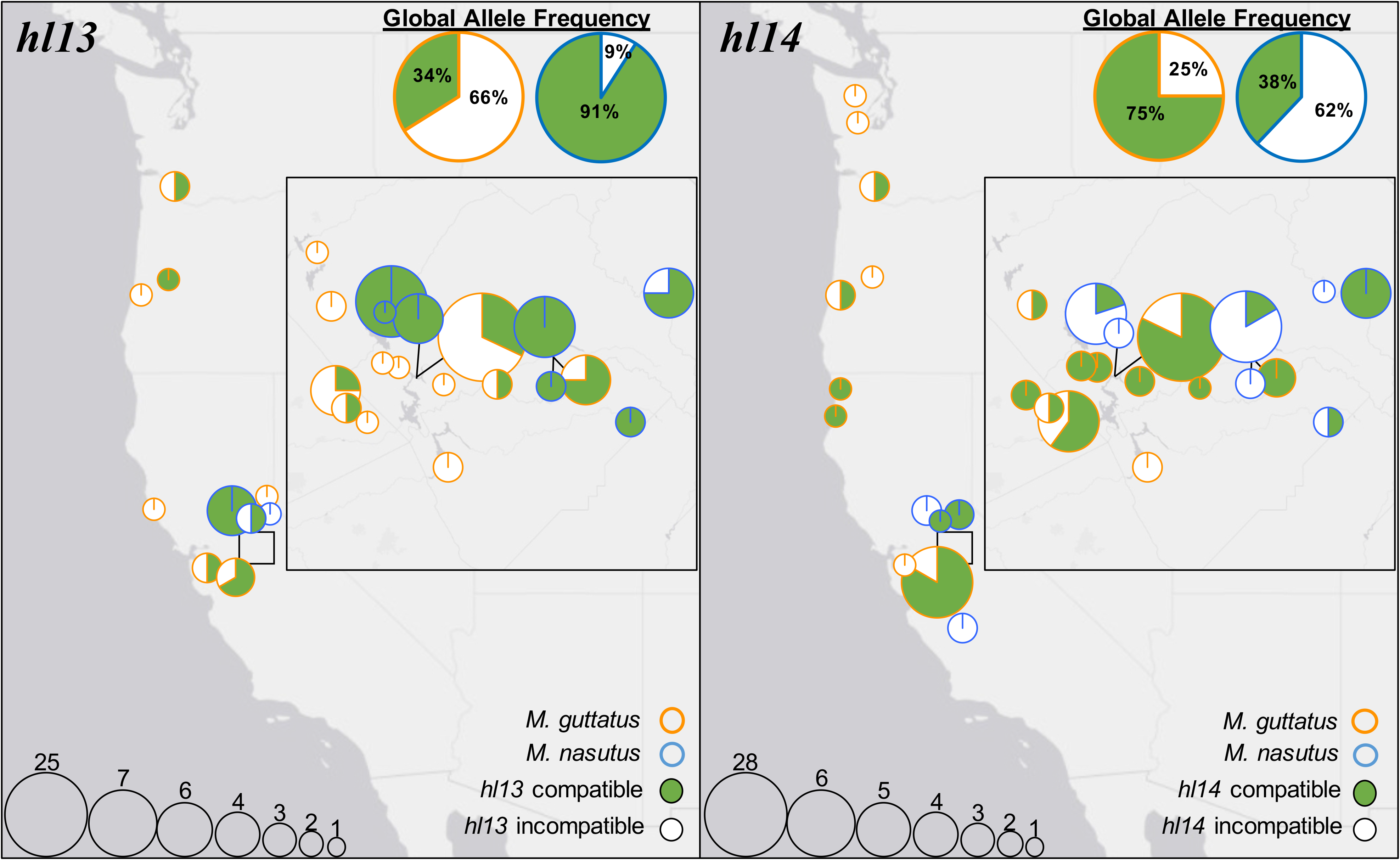
Distribution and frequency of hybrid lethality alleles in *Mimulus.* Circles represent populations sampled across the western United States. Circle size corresponds to the number of alleles tested for each HI locus in a given population (key in bottom left), outer color represents species, and inner color indicates whether alleles were compatible or incompatible (key in bottom right). Global allele frequencies for *M. guttatus* and *M. nasutus* are given in upper right corner. Sympatric populations are connected by black lines.

Patterns of variation at *hl13* and *hl14* within and between *Mimulus* species indicate that hybrid lethality alleles may often come into contact in natural populations. Indeed, we found that *hl13* and *hl14* incompatibility alleles co-occur at both of the sympatric sites of *M. guttatus* and *M. nasutus,* as well as within 50% of the *M. guttatus* populations for which both loci were surveyed (N = 12, Table S3). These findings suggest that interspecific hybridizations and intraspecific crosses might often result in the expression of *hl13-hl14* lethality in nature.

Consistent with this idea, we discovered white seedlings among the selfed progeny of 30% of wild *M. guttatus* individuals tested (N = 17, Table S3).

### Evidence for introgression at hybrid lethality loci

As a first step toward investigating the source of polymorphism at *hl13* and *hl14,* we examined patterns of natural variation in genomic regions surrounding the two loci. We reasoned that if polymorphism is due, at least in part, to recent introgression, we should observe a distinct pattern of linked variation than if it is due only to incomplete lineage sorting (the former is expected to elevate shared variation at linked loci, the latter is not). We surveyed alleles of known incompatibility status *(i.e.,* identified in the crossing experiments described above) for insertion-deletion polymorphisms at linked markers (M236 and M208 for *hl13,* M132 and M241 for *hl14,* Figure S1). Strikingly, at each hybrid lethality locus, we found distinct sets of marker alleles within *M. guttatus* between chromosomes that carried compatible and incompatible variants (Tables 1 and 2, Figure 5). As we detail below, this association between alleles at hybrid lethality loci and linked markers appears to be driven by introgression from *M. nasutus* into *M. guttatus*. Indeed, at each locus, the less common functional variant in *M. guttatus* often carries marker allele combinations that are otherwise confined to *M. nasutus* (or nearly so), indicative of introgression. Within *M. nasutus,* we found little evidence of introgression from *M. guttatus,* but we caution that our sample sizes are prohibitively low (e.g., only three incompatible alleles were sampled at *hl13,* Table 1) and introgression is generally more difficult to detect in this direction (Brandvain et al. 2014).

**Figure 5:**
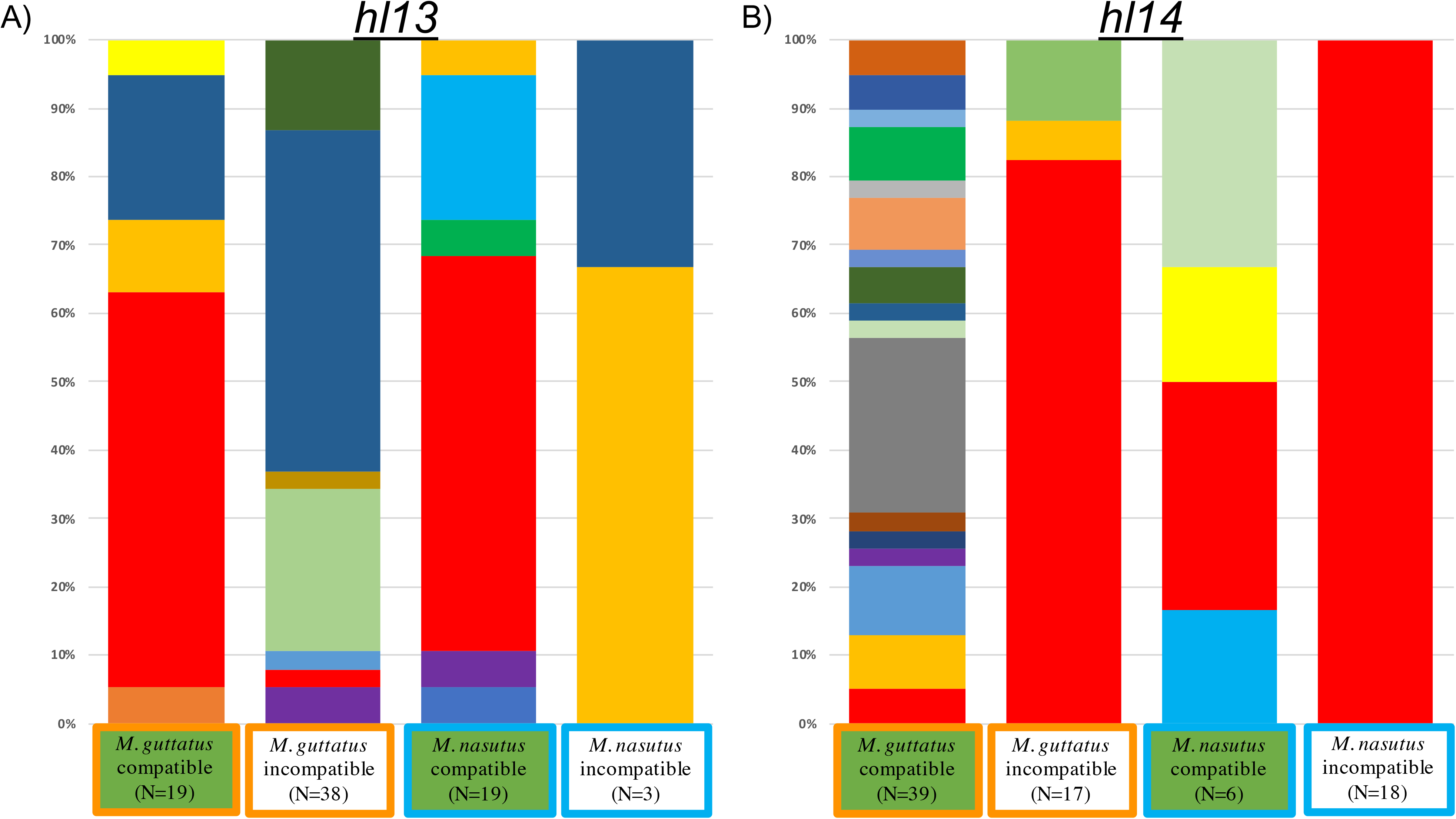
Linked genetic variation at *hl13* and *hl14.* Bars represent linked marker allele combinations associated with compatible and incompatible alleles in *M. guttatus* and *M. nasutus* at (A) *hl13* and (B) *hl14.* Bars are stacked to 100% with sample sizes shown below. Different colors represent distinct linked marker allele combinations generated with markers flanking each locus. Markers M236+M208 and M132+M241 were used to genotype individuals at *hl13* and *hl14,* respectively.

**Table 1.**
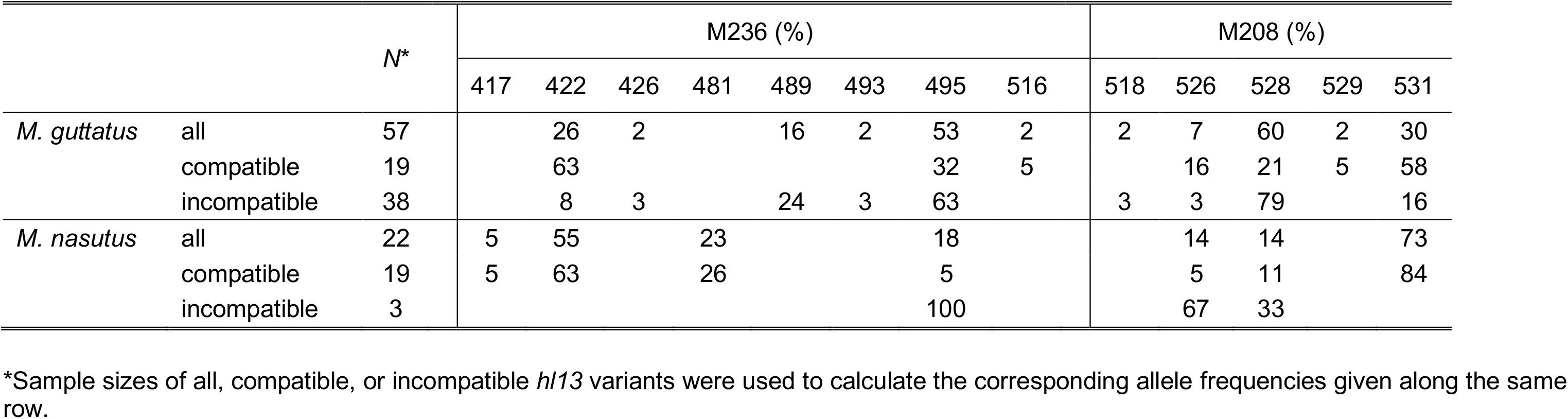
Frequencies of allele sizes at hl13-linked markers M236 and M208 among all, compatible, or incompatible *hl13* variants. Allele frequencies are shown as percentages rounded to the nearest whole number.

Within *M. guttatus,* the two functional classes of *hl13* (compatible and incompatible) differed dramatically in their genetic similarity to *M. nasutus* at linked markers (Table 1). The two markers assayed, M208 and M236, are both proximal to *hl13-pTAC14,* located at distances of 3.3 cM and 7.5 cM, respectively (Figure S1). At M208, the most common *M. nasutus* allele (531) was found in 58% of compatible *M. guttatus* variants but only 16% of incompatible variants. Similarly, at M236, the most common *M. nasutus* allele (422) occurred in 63% of compatible but only 8% of incompatible *M. guttatus* variants. Notably, this pattern of shared variation with *M. nasutus* was highly associated between the two markers: all 11 individuals with 531 at M208 *also* carried 422 at M236. In fact, this single two-locus allele combination (531-422, red bars in Figure 5A) was found in the majority of the compatible *hl13* variants we sampled (58% in *M. guttatus* and in *M. nasutus, N* = 19 each, Table S4). In marked contrast, this same combination was rarely found among incompatible *M. guttatus hl13* variants (occurring in only one of 38 samples, Figure 5A and Table S4). Indeed, for *M. guttatus,* the proportion of two-locus marker allele combinations shared with *M. nasutus* differed significantly between compatible and incompatible *hl13* variants *(X^2^* = 5.000, *P* = 0.025, df = 1). Given that linkage disequilibrium decays rapidly within *M. guttatus* (100-1000bp; (Brandvain et al. 2014, Puzey et al. 2017), our finding of shared variation across a sizeable genetic distance (~7.5 cM between M236 and *hl13*) suggests introgression from *M. nasutus* might act as a source of compatible *hl13* alleles in *M. guttatus.* Under a scenario of neutral evolution at *hl13* within *M. guttatus,* natural selection is expected to favor this pattern of interspecific gene flow, which would reduce the number of hybrid lethal combinations.

At *hl14,* we observed a similar pattern of shared variation between species suggesting that incompatible alleles from *M. nasutus* have also introgressed into *M. guttatus* (Table 2). In this genomic region, we assayed variation at markers that flank *hl14-pTAC14:* M241 is located 6.3 cM proximal to the incompatibility gene and M132 is located 4.5 cM distal to it (Figure S1). At M241, the most common *M. nasutus* allele (283) was found in 82% of incompatible *M. guttatus* variants but only 5% of compatible variants. Similarly, at M132, the most common *M. nasutus* allele (500) occurred in 82% of incompatible but only 18% of compatible *M. guttatus* variants. As with *hl13*, variation at these two *hl14*-linked marker loci is not independent: all 14 *M. guttatus hl14* incompatible variants that carried the 283 allele at M241 also carried the 500 allele at M132. This two-locus allele combination (283-500, red bars in Figure 5B), which was carried by *all* incompatible *hl14* variants sampled from *M. nasutus (N* = 18), dominated the sample of incompatible *hl14* alleles from *M. guttatus* (occurring in 82%, *N* = 17). Accordingly, for *M. guttatus*, the proportion of marker allele combinations shared with *M. nasutus* differed significantly between compatible and incompatible alleles (*X^2^* = 30.679, *P* = 0.0001, df = 1). Taken together, these results suggest that, like at *hl13,* introgression from *M. nasutus* contributes to variation at *hl14* in *M. guttatus.* In this case, however, because gene flow introduces *incompatible* alleles from *M. nasutus* (presumably increasing the number of deleterious *hl13-hl14* combinations within *M. guttatus),* natural selection is expected to oppose it.

**Table 2.** Frequencies of allele sizes at hl14-linked markers M241 and M132 among all, compatible, or incompatible *hl14* variants. Allele frequencies are shown as percentages rounded to the nearest whole number.

|  |  |  | M241 (%) |  |  |  |  |  |  |  | M132 (%) |  |  |  |  |  |  |  |  |  |
| --- | --- | --- | --- | --- | --- | --- | --- | --- | --- | --- | --- | --- | --- | --- | --- | --- | --- | --- | --- | --- |
|  |  |  | 283 | 285 | 287 | 289 | 291 | 292 | 295 | 296 | 406 | 457 | 465 | 469 | 483 | 485 | 490 | 496 | 500 | 503 |
| *M. guttatus* | all | 56 | 29 | 7 | 9 | 30 | 7 | 7 | 4 | 7 | 2 | 2 | 23 | 5 | 7 | 16 | 4 | 2 | 38 | 2 |
|  | compatible | 39 | 5 | 8 | 13 | 44 | 10 | 10 |  | 10 | 3 | 3 | 33 | 3 | 10 | 21 | 5 | 3 | 18 | 3 |
|  | incompatible | 17 | 82 | 6 |  |  |  |  | 12 |  |  |  |  | 12 |  | 6 |  |  | 82 |  |
| *M. nasutus* | all | 24 | 88 | 4 |  | 8 |  |  |  |  |  |  |  | 13 | 4 |  |  |  | 83 |  |
|  | compatible | 6 | 50 | 17 |  | 33 |  |  |  |  |  |  |  | 50 | 17 |  |  |  | 33 |  |
|  | incompatible | 18 | 100 |  |  |  |  |  |  |  |  |  |  |  |  |  |  |  | 100 |  |

\*Sample sizes of all, compatible, or incompatible *h/14* variants were used to calculate the corresponding allele frequencies given along the same row.

## DISCUSSION

Determining whether reproductive isolation is sufficient to prevent hybridizing species from collapsing into a single lineage is a major goal of speciation research. A strong theoretical framework exists for predicting the maintenance of hybrid incompatibilities in sympatry (Gavrilets 1997, Kondrashov 2003, Lemmon and Kirkpatrick 2006, Feder and Nosil 2009, Bank et al. 2012), but there have been few empirical studies investigating these predictions in naturally hybridizing species. With this study, we have begun to fill this gap by determining the geographic distributions and frequencies of incompatibility alleles at two loci that cause hybrid lethality between closely related species of yellow monkeyflower. For each species, we discovered a much higher frequency of incompatibility alleles at one of the two hybrid lethality loci: *hl13* in *M. guttatus* and *hl14* in *M. nasutus.* The implication of this finding is that most naturally formed hybrids will likely carry incompatibility alleles at *both* loci, with the potential to produce lethal offspring. Additionally, because both *hl13* and *hl14* are polymorphic within species, even intraspecific crosses might regularly result in incompatible allele pairings. By examining patterns of genetic variation linked to *hl13* and *hl14,* we show that introgression is a major source of this polymorphism within species, and provide preliminary evidence that natural selection acts to purge incompatible combinations.

### *Incompatibility alleles at* hl13 *and* hl14 *are common and geographically widespread*

In our original genetic characterization of the *hl13-hl14* incompatibility (Zuellig and Sweigart 2018), we used inbred lines derived from the DPR sympatric population of *M. guttatus* and *M. nasutus.* Given this fact, along with our previous discovery of interspecific gene flow at DPR (Brandvain et al. 2014), we suspected that incompatible *hl13-hl14* allele combinations might often arise at this site. Indeed, we found that population-level variation at DPR was well represented by the original inbred lines (Figure 4). Like DPR102-gutt, most *M. guttatus* alleles sampled were incompatible at *hl13* (68%, *N* = 25) and compatible at *hl14* (82%, *N* = 28). Similarly, like DPR104-nas, all *M. nasutus* alleles sampled at DPR were compatible at *hl13 (N* = 4) and incompatible at *hl14 (N* = 2). This pattern of variation means that DPR hybrids will usually be doubly heterozygous at *hl13* and *hl14.* Because both hybrid incompatibility alleles are recessive, the frequency of hybrid lethality will depend on the extent to which F1 hybrids self-fertilize, cross with other hybrids, or backcross with parental species (hybrid lethality will appear in 1/16 of selfed or hybrid intercross progeny, but will not be expressed in most first-generation backcrosses). Of course, at DPR, the lethality phenotype might also be expressed from crosses within *M. guttatus* because this species segregates for incompatible alleles at both *hl13* and *hl14* (Figure 4, Table S3).

Beyond the DPR site, incompatibility alleles at *hl13* and *hl14* are widespread across the ranges of both *Mimulus* species. As a result, in areas of secondary contact, hybrid lethality might be a common outcome of interspecific gene flow between *M. guttatus* and *M. nasutus.* This pattern differs sharply from what has been seen for a two-locus hybrid sterility system between these same species (Sweigart et al. 2006), which likely has limited potential for expression due to the highly restricted distribution of the *M. guttatus* incompatibility allele (Sweigart et al. 2007, Sweigart and Flagel 2015). Additionally, our finding that both *hl13* and *hl14* are polymorphic in roughly half of (apparently) allopatric *M. guttatus* populations means that the lethal phenotype might often contribute to inbreeding depression within *M. guttatus*. In line with this expectation, we observed chlorotic lethal seedlings in the selfed progeny of five wild-collected *M. guttatus* from four populations near DPR (representing 30% of the individuals and 50% of the populations tested, see Table S3). Historically, several studies of inbreeding depression in *M. guttatus* also revealed chlorophyll-deficient lethals (via selfing wild-collected individuals) in multiple populations across the species range (Kiang and Libby 1972, Willis 1992, MacNair 1993). In fact, at the Schneider Creek population, which is roughly 300 km north of DPR, chlorotic seedling lethality was shown to result from recessive alleles at two loci (Willis 1992, MacNair 1993). These same two loci were also implicated in seedling chlorosis segregating in a non-native British population thought to have been introduced from northern California *M. guttatus* (MacNair 1993). Although the causal loci for seedling lethality have not been mapped in these populations, their mode of action and allele frequencies are entirely consistent with what we have observed for the *hl13-hl14* incompatibility. Potentially, then, these two loci have far-reaching impacts on variation in seedling viability within and between *Mimulus* species across their ranges.

### *Origin and maintenance of polymorphism at* hl13 *and* hl14

A major finding of this study is that each *Mimulus* species is polymorphic for both hybrid lethality loci. Functional alleles (incompatible and compatible) show no geographic structure at either locus (in either species) and are often at intermediate frequencies within populations (Figure 4). So, what is the source of this polymorphism at *hl13* and *hl14?* Is it due to incomplete linage sorting, new mutations, or introgression of previously fixed alleles? Given the recentness of species divergence (~200 kya) and extensive natural variation in *M. guttatus* (Brandvain et al. 2014, Puzey et al. 2017), it is easy to imagine that ancestral polymorphism might exist at these loci. In our original genetic characterization of the *hl13-hl14* incompatibility, we showed that hybrid chlorosis occurred in seedlings that inherited *M. guttatus* alleles at *hl13,* which carried nonfunctional versions of the ancestral copy of *pTAC14,* along with *M. nasutus* alleles at *hl14,* which lacked the duplicated copy of *pTAC14* altogether (Zuellig and Sweigart 2018). We do not yet know the evolutionary timing of the *pTAC14* duplication (onto *hl14)* or of the mutation that knocks out function of the ancestral copy on *hl13.* If the duplication arose in ancestral populations and did not go to fixation, presence/absence of *pTAC14* at *hl14* might explain polymorphism for hybrid lethality in descendant lineages of *M. nasutus* and *M. guttatus.* At *hl13,* we already know that the lethality-causing null mutation in the ancestral copy of *pTAC14* is not fixed in *M. guttatus*; the reference genome strain (IM62) carries a copy without the 1-bp insertion and is expressed (Zuellig and Sweigart 2018). Although it is not yet known whether any of these variants has a selective advantage, it seems reasonable to expect that at least some of the polymorphism we observe at *hl13* and *hl14* might be due to mutations that have not yet been fixed by genetic drift.

Strikingly, though, it appears that introgression is a primary driver of polymorphism at both *Mimulus* hybrid lethality loci. This result is not entirely unsurprising because hybridization (particularly introgression from *M. nasutus* to *M. guttatus)* can be substantial in regions of secondary sympatry (Kenney and Sweigart 2016). Many of the *M. guttatus* populations we sampled occur in close proximity to *M. nasutus* and even for those that appear allopatric, it is difficult to rule out sympatry without repeated visits to the site within and across years. However, although introgression between these species is generally plausible, we did observe *M. nasutus*-like variants in several *M. guttatus* individuals from what seem to be genuinely allopatric populations. One possibility is that the *hl13-* and hl14-linked markers do not always accurately diagnose introgression across these large chromosomal fragments. Indeed, recombination in the wild parents might have disassociated marker and incompatibility alleles (see Methods), and marker homoplasy (e.g., distinct indels of the same size) could give a false signal. Going forward, a promising approach will be to identify distinct *pTAC14* variants by direct sequencing and examine patterns of linked genomic variation at a much finer scale (e.g., (Brandvain et al. 2014, Kenney and Sweigart 2016).

Of course, a key question is which forces maintain polymorphism at these hybrid incompatibility loci in natural populations. Theory predicts that hybrid incompatibility alleles evolving neutrally within species cannot be maintained in the face of interspecific gene flow because it reveals their deleterious effects in hybrids (Gavrilets 1997, Kondrashov 2003, Bank et al. 2012, Muir and Hahn 2014). In this system, because of asymmetric gene flow from *M. nasutus* into *M. guttatus* (Sweigart and Willis 2003, Brandvain et al. 2014), natural selection is expected to favor introgression of the *M. nasutus* allele at *hl13 (i.e.,* the functional, ancestral copy of *pTAC14)* into *M. guttatus,* effectively disabling the *hl13-hl14* incompatibility. Consistent with this idea, many of the compatible *hl13* variants we found in *M. guttatus* are embedded in large blocks of *M. nasutus-like* variation, indicative of recent introgression events. Even more intriguing is the finding that several other compatible *hl13* variants carried marker alleles typical of *M. guttatus*, suggesting that selection might have preserved introgressed *M. nasutus* alleles long enough for recombination to erode the signal of gene flow (although we cannot rule out ancestral polymorphism for these individuals). At *hl14,* where we expect natural selection to oppose introgression from *M. nasutus* into *M. guttatus,* polymorphism seems to be driven almost exclusively by recent hybridization, suggesting that incompatible alleles are purged quickly *(i.e.,* they do not persist long enough to recombine with linked markers). This localized effect at *hl14* is consistent with genome-wide patterns of variation in this system (Brandvain et al. 2014, Kenney and Sweigart 2016) and others (Payseur et al. 2004, Teeter et al. 2008, Schumer et al. 2018) that indicate pervasive selection against introgression. Under such a scenario, polymorphism at *hl13* and *hl14* is expected to be transitory: DPR and other sympatric populations are presumably on their way to fixing compatible alleles.

An alternative possibility is that incompatibility alleles are stably maintained within populations. Based on their molecular lesions, we have speculated that the *hl13* and *hl14* incompatibility alleles are selectively neutral within species (one is not expressed, one is missing: see Zuellig and Sweigart 2018). However, if these alleles are advantageous within species (e.g., because knocking out one copy maintains proper dosage), intermediate frequencies might reflect a balance between their positive effects within species and negative effects in hybrids. Additionally, without knowing exactly how much gene flow is occurring, it is difficult to determine if deleterious two-locus combinations occur frequently enough for selection to remove hybrid lethal alleles from the population. Furthermore, if F1 hybrids usually backcross to *M. guttatus*, incompatible alleles from *M. nasutus* will almost always be shielded in heterozygotes. Because selection against incompatibility alleles occurs only in the double homozygote, the conditions for maintaining both alleles through migration selection balance might not be particularly restrictive. This situation is similar to the equilibrium described by (Fisher 1935) for maintaining lethal alleles at duplicate loci. In his discussion of tetraploidy, he pointed out that for an infinite population size with equal mutation rates at the two loci, the frequency of lethal double homozygotes will equal the mutation rate. As a result, there can be many different combinations of lethal allele frequencies at the two loci (from equal to highly asymmetric) that will maintain mutation-selection balance. Of course, for the highly selfing *M. nasutus*, we never found both incompatibility alleles within populations or white seedlings among selfed progeny, consistent with the idea that incompatibility alleles are purged immediately when made homozygous.

With the genes for *hl13* and *hl14* now identified, a major goal of our future studies will be to pinpoint the evolutionary processes that maintain variation for this hybrid incompatibility in nature. Because this *Mimulus* system is so tractable for field studies, it will be possible to follow the frequencies of specific *pTAC14* alleles within and across years, and to determine the fitness effects of incompatibility alleles within species and in hybrids. Such information will be critical for predicting whether this hybrid incompatibility will be stably maintained or dissolve with interspecific gene flow, and for understanding its role (if any) in *Mimulus* speciation.

---

**Supplemental Figure 1**: Genetic distance between size-polymorphic markers and *pTAC14* duplicates at *hl13* and *hl14.* Genetic distances were calculated from a F2 mapping population of white F2 seedlings (DPR104-nas x DPR102-gutt) described in Zuellig and Sweigart (2018).

## AUTHOR CONTRIBUTIONS

M.P.Z. and A.L.S. designed research; M.P.Z. and A.L.S. performed research; M.P.Z. and A.L.S. analyzed data; M.P.Z. and A.L.S. wrote the manuscript.

## ACKNOWLEDGEMENTS

We thank Sam Mantel, Brad Whitfield, Fernando Fernandez, Taylor Harrell, Ananya Moorthy, Rachel Hughes and Rachel Kerwin for their help in the greenhouse and lab. Dave Hall, CJ Tsai, Mike Arnold, John Burke, and Kelly Dyer provided valuable advice throughout the study and helpful suggestions on earlier drafts. We also thank members of the Sweigart, Bergman, Dyer, Hall, and White labs for providing thoughtful comments on a draft of this manuscript, which improved its quality. This work was supported by a National Institute of Health T32 Fellowship (GM007103) to M.P.Z. and a National Science Foundation grant (DEB-1350935) to A.L.S.

